# Exploring the Mechanism of Olfactory Recognition at the Initial Stage by Modeling the Emission Spectrum of Electron Transfer

**DOI:** 10.1101/641738

**Authors:** Shu Liu, Rao Fu, Guangwu Li

**Affiliations:** Department of Anatomy, Anhui Medical University, Meishan Road 81, Hefei 230032, China; Department of Anesthesiology, New Jersey Medical School, Rutgers, the State Univerisity of New Jersey, 185 South Orange Avenue, Newark, New Jersy, 07103

**Keywords:** Olfaction, Electron Transfer, Huang–Rhys factor, Emission Spectrum, Isotope Effect

## Abstract

Olfactory sense remains elusive regarding the primary reception mechanism. Some studies suggest that olfaction is a spectral sense, the olfactory event is triggered by electron transfer (ET) across the odorants at the active sites of odorant receptors (ORs). Herein we present a Donor-Bridge-Acceptor model, proposing that the ET process can be viewed as an electron hopping from the donor molecule to the odorant molecule (Bridge), then hopping off to the acceptor molecule, making the electronic state of the odorant molecule change along with vibrations (vibronic transition). The odorant specific parameter, Huang–Rhys factor can be derived from *ab initio* calculations, which make the simulation of ET spectra achievable. In this study, we revealed that the emission spectra (after Gaussian convolution) can be acted as odor characteristic spectra. Using the emission spectrum of ET, we were able to reasonably interpret the similar bitter-almond odors among hydrogen cyanide, benzaldehyde and nitrobenzene. In terms of isotope effects, we succeeded in explaining why subjects can easily distinguish cyclopentadecanone from its fully deuterated analogue cyclopentadecanone-d28 but not distinguishing acetophenone from acetophenone-d8.

## Introduction

The sense of smell (olfaction) is vital to the survival. A great progress has been made in understanding the physiological and biochemical basis of olfaction, but it remains elusive regarding the primary reception mechanism at the initial stage. One argument is associated with the “lock and key” theory[1-4], it has been widely believed that odorant molecules bind to specific receptors through conventional molecular interactions, leading to a conformational change in the receptor that activates intracellular signals. However, this theory could not predict the odor character of molecules and rational odorant design due to the large number of ORs and the breadth of OR tuning[5]. An alternative concept is the “vibration” theory[6-10], in which researchers speculate that olfaction is a spectral sense and the olfactory event is triggered by an electron transfer (ET) within the OR[11]. Although the vibrational theory was successfully to be used to predict the smells of certain well-documented odorants[12], it has not been widely accepted due to its failure in multiple instances.

ET is an integral part of many biological processes, such as photosynthesis and respiration[13-18]. Biophysical investigations have produced a strikingly detailed picture of electron transfers occurring between metal-containing cofactors in a metalloprotein[19-22]. Some experimental studies observed that the OR is also a metalloprotein; the transition metal ions (e.g., Zn^2+^, Cu^2+^, and Cu^+^) could coordinate odorant molecules[23,24]. Pshenichnyuk et al. proposed that there exists ET through an odorant molecule in the OR, the electron-accepting properties of odorant molecules are involved in their smell recognition[25]. Brookes et al.[26] proposed that the source of excess electrons is likely a reducing (oxidizing) agent in the cell fluid. Bittner et al.[27] posited that by projecting the impulsive force onto the internal vibrational modes of odorant molecule, only those modes resonant with the tunneling gap will excite impulsively contribute to the inelastic transfer rate. Horsfield et al[28] made an attempt to bring together the various theories into a single formalism regarding the olfactory recognition step.

Once a suitable odorant molecule enters the binding pocket of the receptor and docks successfully, i.e., the shape fit and the orientation are correct according to van der Waals and electrostatic interactions, the odorant molecule may act as a bridge (B) molecule between the D and A molecules and forms the so-called Donor–Bridge–Acceptor (DBA) system. If the lowest unoccupied molecular orbital (LUMO) of the B molecule is approximately resonant with the LUMOs of the D and A molecules, an extra electron may jump from the D molecule to the B molecule, where it stays for a while before hopping off to the A molecule[29]. The odorant molecule (B) simultaneously changes its electronic state with an excitation of one or more phonons; we refer to this combination of vibrational and electronic transitions as the vibronic transitions. As illustrated in Figure 1, during the electron transfer process, the odorant molecule undergoes four vibronic states, S0 NEU→Sυ ANI→S0 ANI→Sυ NEU, representing a neutral state with a ground vibrational state, anionic state with vibrational excitation, anionic state with a ground vibrational state and neutral state with vibrational excitation, respectively. With electron absorption, the odorant molecule starts in the vibronic state S0 NEU and finish in Sυ ANI; with electron emission, i.e., an extra electron jump from the odorant molecule (B) to A molecule, the B molecule starts in S0 ANI and finish in Sυ NEU.

**Figure 1.**
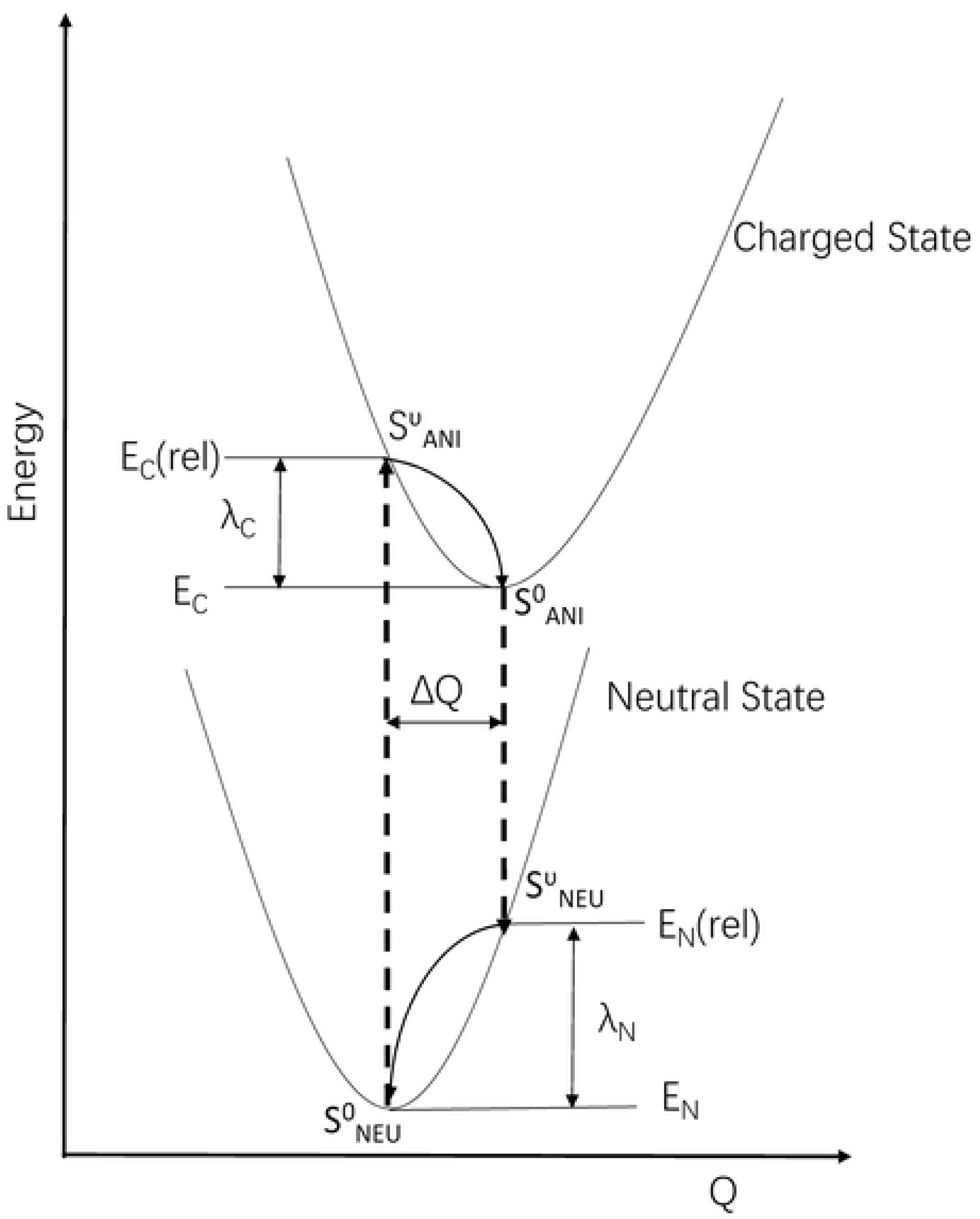
Sketch of the potential energy surfaces for the neutral state and charged state, showing the four vibronic states, S0 NEU, Sυ ANI, S0 ANI and Sυ NEU, the vertical transitions (dashed lines), the normal mode displacement ΔQ, and the relaxation energies λ_N_ and λ_C_.

Up to now, no OR has been crystallized yet, it is unknown where the electron donor (D) site and acceptor (A) site are within the OR, some parameters (the hopping integral between donor and acceptor states, the free energy and the intermolecular reorganization energy) of ET rates possess a higher degree of uncertainty, it is difficult to investigate what conditions lead to high ET rates, and what do not. For the weak interactions (van der Waals and electrostatic interactions) between the B molecule and the D and A molecules, we consider that the electronic coupling of the odorant molecule to the D (A) molecule is unlikely to modify the free molecule properties. Therefore, we only need to consider the individual odorant molecule to study the vibronic transition properties of the odorant, without involving the odorant molecule B and the D (A) molecule as a complex entity, so that the specific parameters (the vibrational frequency ω and Huang-Rhys factor 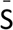) of ET rates can be derived from ab initio calculations.

Huang–Rhys factor 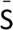 provides a quantitative description of electron-vibrational coupling property. For the 0−υ vibronic transition, the intensity I0−υ j in each vibrational mode j can be expressed by a Poisson distribution with the Huang–Rhys factor 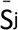 as an argument[30,31]:

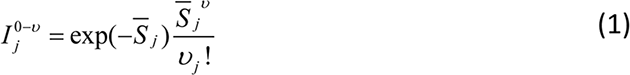

Where υ_j_ denotes the final vibrational quantum numbers (phonons). If an electron transfer across the odorant within the OR and the vibronic transition occurs, the ET spectra of the odorants (after convolution) should correlate with their odor character. The band number, width and position of ET spectrum may be responsible for molecule’s odor character. Via extensive comparative spectral analysis, we revealed that the emission spectrum of ET could act as an objective evaluation criterion for odor character.

Two different tests were used to validate correlation between the emission spectrum of ET and the odor character of molecules as following described: (i) To explore why some odorants have different structures but possess similar smells, such as hydrogen cyanide (HCN), benzaldehyde and nitrobenzene. To be noted, the reason why HCN possesses bitter almond odor character has not yet been reasonably interpreted by “lock and key” theory. We simulated the ET spectra of these odorants, finding that the emission spectra were highly overlapped at a certain vibrational frequency. (ii) Isotope effects on odor character have been controversial between supporters and opponents of vibrational theory. The experimental studies of Gane et al.[32] showed that subjects are incapable of distinguishing acetophenone and its fully deuterated analogue acetophenone-d8 but can easily distinguish between cyclopentadecanone and cyclopentadecanone-d28. In this study, we were able to perfectly interpret this previous experimental results by comparing the ET emission spectrum of one molecule with its deuterated analogue.

## Methods

All quantum chemical calculations were performed using Gaussian 09[33] and analyzed using Multiwfn[34]. For seven odorant molecules of interest (HCN, benzaldehyde, nitrobenzene, acetophenone, acetophenone-d8, cyclopentadecanone, and cyclopentadecanone-d28), we first perform geometry optimizations of the neutral states using the B3LYP density functional[35-37] with the 6-31G basis set[38,39] in a vacuum; the same functional and basis sets were then used to perform geometry optimizations of the anionic states at the neutral equilibrium geometry. Subsequently, single-point calculations were performed using a more flexible basis set augmented with diffuse functions (6-311+G**)[40,41]. IR spectra were generated by application of a broadening factor of 4 Hz and compiled for presentation using Excel. The intramolecular reorganization energies and the Huang-Rhys factors 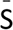were computed using a modified version of the DUSHIN program developed by Reimers[42].

### Calculation of the intramolecular reorganization energy and Huang–Rhys factor

The reorganization energy λ represents the changes to the nuclear positions corresponding to the charge movement; the total reorganization energy λ includes the intramolecular reorganization energy (λ_i_) and the external reorganization energy (λ_S_) components. The intramolecular reorganization energies λ_i_ describe the strength of the electron-vibration coupling energy of a charge localized on a single molecule and can be estimated in two ways, with an adiabatic process using adiabatic potential energy surface (AP) and with a normal mode (NM) analysis. In the latter procedure, the λ_i_ of the molecules were decomposed into the normal-mode contributions, and the contribution of each vibrational mode to λ_N(C)_ can be obtained by expanding the potential energies of the neutral and charged states in a power series of the normal coordinates based on harmonic oscillation approximation[43-48]:

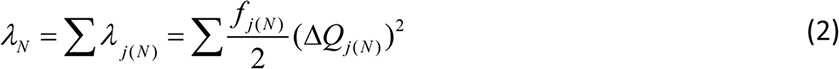

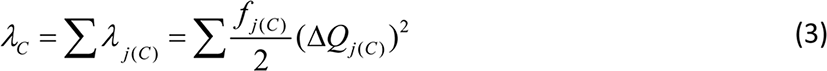

Here, ΔQ_j(N)_ and ΔQ_j(C)_ are the change in equilibrium geometry between the neutral state and the charged state along the *j*-th normal coordinate; f_j(N)_ and f_j(C)_ are the corresponding force constants of the neutral and charged states.

The Huang–Rhys factor 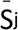 corresponds to the intramolecular reorganization energy λ_j_ and the corresponding vibrational energy ħω_j_:

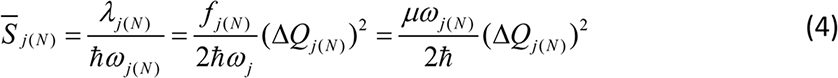

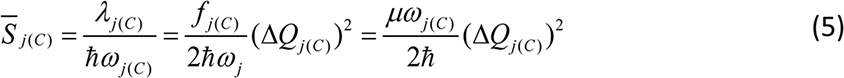

where ω_j(N)_ and ω_j(C)_ are the effective vibrational frequencies in the neutral and charged states, respectively, and μ is the reduced mass for a normal mode of vibration.

The normal modes of one electronic state are no longer orthogonal to the normal modes of the other electronic state, generally called the the Duschinsky rotation effect[49-51]. The displaced and mixed modes can be determined by Duschinsky’s transformation, expressed by Eq. (6)[50]:

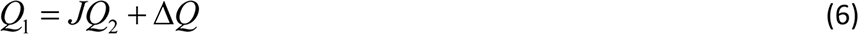

Here, Q_1_ and Q_2_ represent the normal coordinates of the electronic state S_NEU_ and S_ANI_ for absorption transition, or the normal coordinates of the electronic state S_ANI_ and S_NEU_ for emission transition, respectively. ΔQ is a vector of the displacement, and J is the Duschinsky rotation matrix.

The overall line shape is assumed as a convolution of the intensity distribution function with the δ-function replaced by a line shape function G_N_(ω)[52,53]. Jortner[54] proposed that for the ET rate expression, the contribution of the solvent modes was lumped into a Gaussian line shape. In this study, the calculated spectral intensities are a set of vertical lines underlying vibrational modes (abscissa). Following Turin’s suggestion[11] that the resolution of human olfactory spectrometers is approximately 400 cm^-1^ for the width of inelastic electron tunneling spectroscopy (IETS), we applied a Gaussian function with a standard deviation of 100 cm^−1^ to convolute the vertical lines. In this way, the simulated spectrum became a smooth curve with characteristic of width and intensity, which was similar to the experimental spectrum, thus making it convenient to compare different molecules.

## Results

### Spectral Comparison of HCN, Benzaldehyde and Nitrobenzene

The results of Huang-Rhys factors 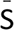 for HCN, benzaldehyde and nitrobenzene are listed in Table 1, Table S1 and Table S2. When an exogenous electron is added, the electronic state of HCN is changed from a neutral state to an anionic state, the spin population is primarily distributed at the C≡N bond (Table S3), the geometry changes correspond to an elongation of these three bonds, which lead to the effect on intramolecular reorganization energy, the largest at 2149 cm^−1^ in the neutral state and 1704 cm^−1^ in the anionic state, and the corresponding Huang-Rhys factors 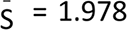 and 1.594, respectively. During the vibronic transition, the HCN bending and C-H bond length will be more or less unchanged, their Huang-Rhys factors 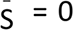 and 0.084 are in the neutral state, and 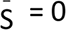 and 0.029 in the anionic state (Table 1).

**Table 1.** The Huang-Rhys Factors 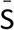 and intramolecular reorganization energies λ_i_ (eV) for each vibrational frequency ω_i_ (cm^-1^) of HCN in its neutral and anionic states.

| Neutral |  |  | Anionic |  |  | vibrational mode |
| --- | --- | --- | --- | --- | --- | --- |
| $\omega_i$ | $\bar{S}$ | $\lambda_i$ | $\omega_i$ | $\bar{S}$ | $\lambda_i$ | |
| 778 | 0 | 0 | 696 | 0 | 0 | bend |
| 2149 | 1.978 | 0.527 | 1704 | 1.594 | 0.337 | C≡N stretch |
| 3500 | 0.084 | 0.036 | 3446 | 0.029 | 0.013 | C-H stretch |

According to Eq. (1), we can simulate the ET spectra with the Huang-Rhys factors 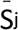 and the final vibrational quantum numbers υ_j_. For the ET reaction in the mixed quantum classical regime, υ_j_ depends on the magnitude of the 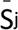. Jortner[55] has reported that for an ET process at high temperature the activation energy exhibits a parabolic dependence, 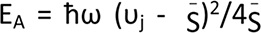, for activationless processes 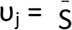. Therefore, based on the Huang-Rhys factors 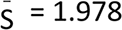 and 1.594 for HCN, we speculated that the vibronic transition excites two phonons in this odorant. Figure 2A shows that the calculated ET absorption and emission spectra of HCN before (vertical lines) and after (smooth curve) convolution; the emission band (approximately 1704 cm^-1^) is left-shifted compared with the absorption band (approximately 2149 cm^-1^).

**Figure 2.**
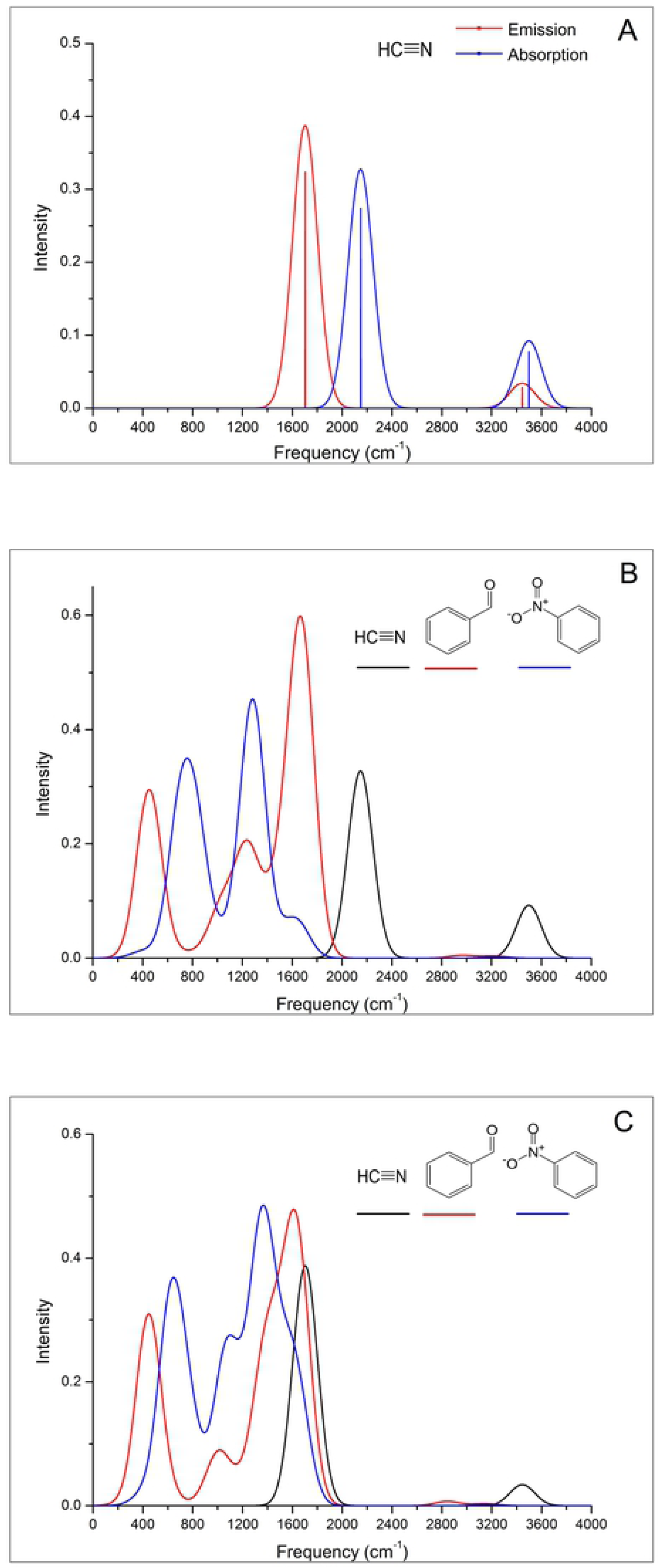
A. The structure and ET absorption and emission spectra of HCN. The calculated peaks are shown before (vertical bars) and after (smooth curve) convolution with a Gaussian (s.d.=100 wavenumbers). B. The structures and ET absorption spectra of HCN, benzaldehyde and nitrobenzene. C. The structures and ET emission spectra of HCN, benzaldehyde and nitrobenzene.

It should be noted that the ET emission spectra are not the vibrational spectra of the negatively charged states, similarly, ET absorption spectra are not the vibrational spectra of neutral states, because ET spectra origin from vibronic transition and are calculated from Huang-Rhys Factors 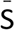. For HCN, the intensity value of the ET spectrum of HCN is not directly related to the IR intensity of the corresponding vibrations, as illustrated in Figure 2A and Figure S1A, the C-H stretching mode and bending mode have the large IR intensity, but the intensity of the ET spectrum is very low. For the C≡N stretching mode, the case is the opposite. The reason can be explained as follows: for the IR of HCN in the ground state, the C≡N stretching does not produce a change in the dipole moment, and the moment remains zero throughout the vibration by symmetry, but the C-H stretching mode and bending mode can produce a change in the dipole moment during the vibration, leading to IR activity.

Figure 2B and Figure 2C show that both ET absorption and emission spectra of benzaldehyde are composed of two prominent peaks at approximately 450 cm^-1^ and 1600 cm^-1^, and a lower, smaller peak at 1000 cm^-1^. The first peak in the low-frequency region of the spectra is dominated by a vibronic progression originating from the coupling to vibrational mode with bending vibration in-plane at a calculated 453 cm^-1^ in the neutral state and 448 cm^-1^ in the anionic state. The third prominent peak in the high-frequency region of the spectra is dominated by vibronic progression originating from the coupling vibrational modes are the aromatic C=C stretch as well as C=O stretch at 1660 cm^-1^ and 1707 cm^-1^ in the neutral state and at 1578 cm^-1^ and 1654 cm^-1^ in the anionic state (Table S1).

Figure 2B and Figure 2C also show that both the ET absorption and emission spectra of nitrobenzene are composed of two peaks. For the first peak of the absorption spectrum, the obvious vibronic feature corresponding to the vibrational mode with benzene ring bending vibration in-plane is at a calculated 699 cm^-1^, the nitro group scissoring vibration is at a calculated 831 cm^-1^ in the neutral state. For the first peak of the emission spectrum, the coupling vibrational modes are calculated at 626 cm^-1^ and 772 cm^-1^ in the anionic state. The second peak of the absorption spectrum, which is in the high-frequency region and is primarily dominated by vibronic progression originating from the coupling to vibrational mode with the benzene ring C-H rocking vibration in-plane and the N=O stretch in the nitro group, is calculated at 1290 cm^-1^. For the second peak of the emission spectrum, the coupling vibrational modes are calculated at 1069 cm^-1^, 1359 cm^-1^, 1524 cm^-1^ and 1641 cm^-1^ in the anionic state, respectively (Table S2).

Quantitative spectral comparison could be used to investigate which of the two types of ET spectrum (absorption and emission) could be the odorant characteristic spectrum. As shown in Figure 2B, comparing the calculated ET absorption spectrum of HCN with that of benzaldehyde and nitrobenzene using an overlap criterion, the prominent peak at approximately 2149 cm^-1^ of HCN has no overlap with the peaks of the other two odorants. In contrast, compared with the emission spectra, as shown in Figure 2C, the prominent peak of HCN at approximately 1704 cm^-1^ has 90% band overlap with the third peak of the benzaldehyde, nitrobenzene also gives a good fit to the HCN band, the second peak of nitrobenzene considerably overlaps with the third peak of benzaldehyde.

Remarkably, for the third peak of the emission spectra of benzaldehyde, the coupling vibrational modes are the aromatic C=C stretch as well as C=O stretch at 1578 cm^-1^ and 1654 cm^-1^ in the anionic state, which involve the odotope (C=C–C=O) identifies by Zakarya et al. [56] as responsible for the odor of bitter almonds. We assume that the molecules with similar odors should have similar spectra, the common spectral feature for odorants correlates with a certain odor character. Because the three structurally different bitter-almond molecules have a common spectral feature at approximately 1600 cm^-1^ in the emission spectra, we deduced that the emission spectrum of ET can act as the odorant characteristic spectrum.

To verify the hypothesis that the ET emission spectrum can act as the odorant characteristic spectrum, we expanded the scope of test examples and chose a broad set of odorants belonging to widely different structural and odor classes, such as musks, ambers, woods, violets, and some of the other bitter almonds. The spectral comparison results of these odorants are consistent with the spectral comparison of the small subset (HCN, benzaldehyde and nitrobenzene). For odorants with similar odors, the common spectral feature can be found in the emission spectra, no matter whether their structures are the same, while in the absorption spectra, we did not find any common features.

Some remaining discrepancy could be due to Duschinsky mixing, because we adopt the DUSHIN program[42] to calculate the normal-mode-projected displacements and Duschinsky rotation matrices with the use of curvilinear coordinates, preliminary calculations of Duschinsky matrices point to the presence of some Duschinsky mixing among the vibrations in benzaldehyde and nitrobenzene (Table S4 and S5). Additionally, to allow large-amplitude curvilinear motions to be partitioned into individual vibrational components using the machinery of normal-mode analysis, the DUSHIN program ignores the coordinate dependence of the Jacobian describing the curvilinear-coordinate transform and hence ignores all kinetic energy anharmonicity effects. To further improve the spectral similarity for molecules with the same odor, investigations are currently in progress.

### Isotope effect

A classical approach to test the plausibility of the vibrational theory is to compare the odor character difference between the deuterated isotopomers. However, some psychological experiments can hardly be interpreted by the previous vibrational theories. Keller and Vosshall[57] reported that naive subjects are incapable of distinguishing acetophenone and acetophenone-d8. Gane et al.[32] confirmed their results on trained subjects. In the same article, Gane et al. additionally reported that trained subjects can easily distinguish deuterated and undeuterated musk odorants, cyclopentadecanone and cyclopentadecanone-d28.

In this study, we attempt to interpret the controversial phenomenon of the experimental results by ET spectroscopic analysis. As shown in Figure 3A, both emission spectra of acetophenone and acetophenone-d8 have three peaks after convolution. For the first peak at approximately 400 cm^-1^ in the low-frequency region, both spectra largely consist of a series of vibronic features corresponding to the molecular skeleton vibration. The obvious vibronic feature in non-deuterated corresponds to vibrational mode at a calculated 480 cm^-1^ with CH_3_-CO-C_6_H_5_ scissoring vibration, as well as benzene ring breathing vibration in the anionic state, with a Huang-Rhys factor of 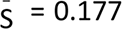; for deuterated acetophenone, the corresponding mode is calculated at 450 cm^-1^ with a Huang-Rhys factor of 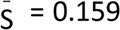 (Table S6 and Table S7). For the second peak, the obvious vibronic feature for non-deuterated acetophenone corresponding to the vibrational mode with benzene ring breathing vibration and sp^3^ C-H deformation vibration is calculated at 1014 cm^-1^, with a Huang-Rhys factor of 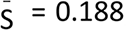; for deuterated acetophenone, the corresponding mode is calculated at 811 cm^-1^ with a Huang-Rhys factor of 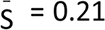. For the third peak, the emission spectra largely consist of a series of vibronic features corresponding to the benzene ring C-H anti-symmetric stretching vibration approximately 1630 cm^-1^ for both non-deuterated and deuterated acetophenone.

**Figure 3.**
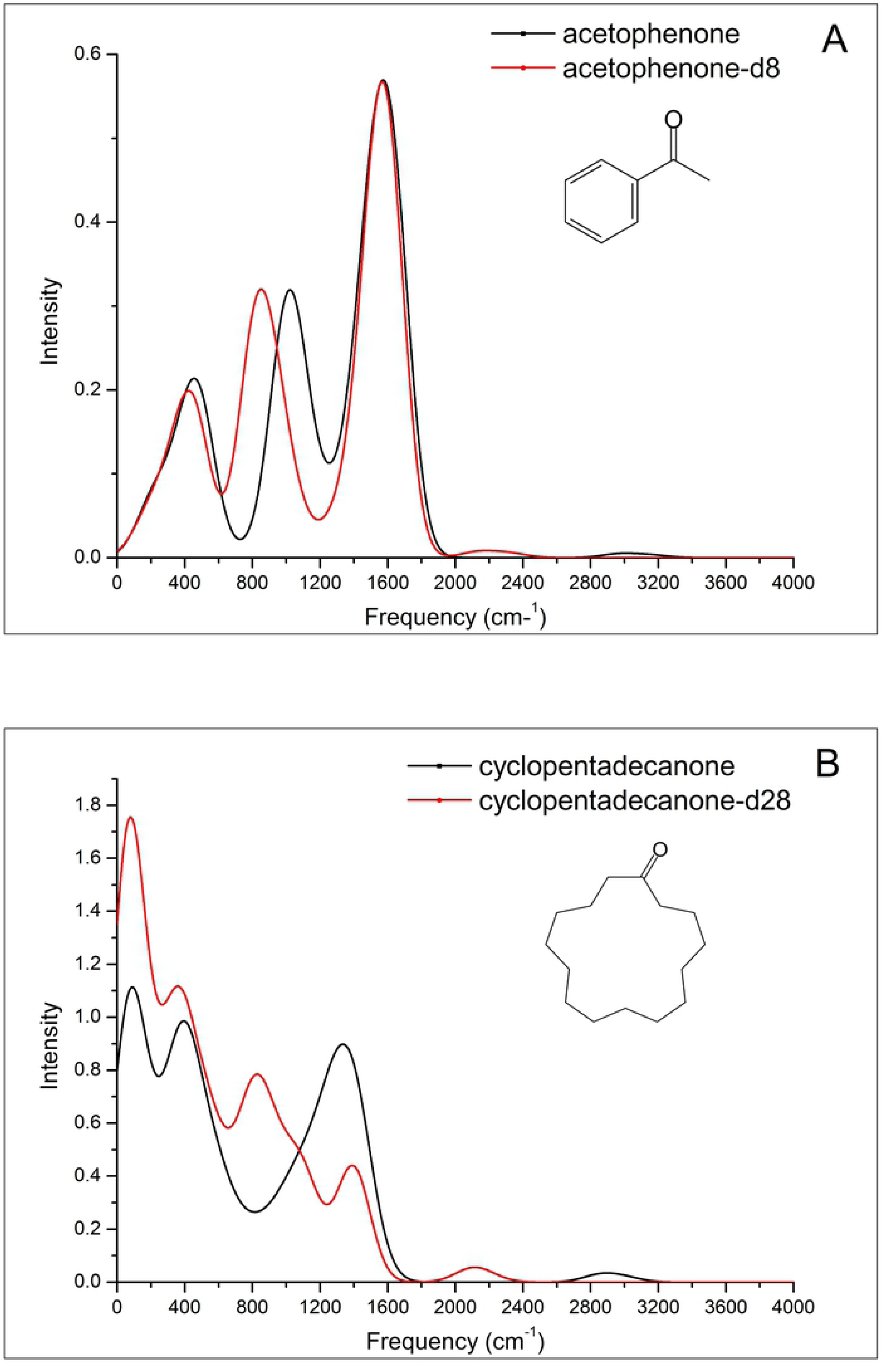
A. The structures of acetophenone and ET emission spectra of acetophenone and acetophenone-d8. B. The structures of cyclopentadecanone and ET emission spectra of cyclopentadecanone and cyclopentadecanone-d28.

By spectroscopic analysis, we can see that the first and third peak of the two isotopes are overlapping at approximately 400 cm^-1^ and 1630 cm^-1^; for the second peak, both the shape and the intensity of the two isotopes are almost identical, but the position of the deuterated peak showed a left-shift of approximately 200 cm^-1^ compared to the non-deuterated peak. According to Turin’s theory[11], human olfactory receptors constitute biological spectrometers with poor resolution (∼400 cm^-1^), it is reasonable to say that ordinary human subjects cannot detect the effect of deuterium on odor character in acetophenone. Nevertheless, the odor character difference between acetophenone isotopomers, which are associated with the slightly different locations of the second peaks of ET emission spectra, could be detected by some animals with keen olfactory sense. For instance, Franco et al.[58] has reported that flies can use olfaction alone to discriminate isotopic acetophenone odorants.

As presented in Figure 3B, cyclopentadecanone has a macrocyclic ketone structure with a 15-membered ring. For its deuterated and undeuterated isotopomers, each molecule has more than 120 vibrational modes (Table S8 and Table S9). The intensities of the vibronic emission spectrum in each vibrational mode may not be large, but they can be combined to form a large peak after convolution. The spectrum of non-deuterated cyclopentadecanone has 3 peaks within 2000 cm^-1^, while the spectrum of cyclopentadecanone-d28 has 4 peaks. The first and second peak of the two isotopes are at the same position, while the intensity of deuterated cyclopentadecanone is greater than that of non-deuterated cyclopentadecanone. Given the emission spectra of the two isotopic cyclopentadecanone odorants are completely different, their odor differences should be easily perceptible.

To interpret deuteration can alter the odor character of cyclopentadecanone but not that of acetophenone, Gane et al.[32] proposed that deuteration exerts the largest effect on the parts of the vibrational spectrum involving C-H motions, odor character might be detectable in odorants containing more C-H groups, since the C-H bond is weakly polar, a bond of low polarity may be difficult to detect by smell. In contrast to acetophenone which contains only 8 carbons and 8 hydrogens, cyclopentadecanone has 15 carbons and 28 hydrogens, more than 3 times the number of vibrational modes involving hydrogens than in acetophenone. This explanation does not satisfy the opponents of the vibrational theory. Hettinger[59] stated that the comparison between C-H with C-D stretching modes showed an obvious isotope shifts from the 3000 cm^-1^ region into the 2200 cm^-1^ region, the difference is as great as half of the entire spectral range in human vision or half an octave in hearing. Accordingly, the odor character difference between acetophenone isotopomers should be easy to detect; however, this is not the case.

As shown in Figure 3A and 3B, there’s a very low, flat peak at the 3000 cm^-1^ for non-deuterated acetophenone and cyclopentadecanone, the corresponding modes are C-H stretch modes in the anionic state; and a similar low, flat peak at the 2200 cm^-1^ region for deuterated acetophenone and cyclopentadecanone, the corresponding modes are C-D stretch modes in the anionic state. By spectroscopic analysis, we conclude that C-H and C-D stretch modes have little effect on the switch from the anionic state to the neutral state geometries of acetophenone and cyclopentadecanone, and have little to do with odor recognition, despite the significant change in C-H stretch frequency upon substitution by deuterium. Both C-H and C-D stretch modes have the large IR intensity, but the corresponding intensity of the ET emission spectra are very low, this proved once again that ET emission spectra are not the vibrational spectra of the negatively charged states.

## Discussion

Charge transfer (CT) is ubiquitous in biology [16-21]. There are two types of CT: (1) The hole transfer (HT) in the case of the radical cations, and (2) The electron transfer (ET) in the case of the radical anions. The type of CT may occur via two different mechanisms: the super-exchange transfer or the sequential (hopping) transfer[60-65]. Correspondingly, we think placing an odorant molecule (B) between the D and A molecules allows the ET to occur either as a single coherent scattering event from D to A directly (super-exchange transfer), or as a sequence of two incoherent hops, first from D to B then from B to A (sequential hopping regime). Brookes et al.[26] has proposed a super-exchange regime that ET occurs from D to A directly within the OR, coupling of a single vibrational frequency of the odorant, and the tunneling electron does not have to go through the molecular orbitals on the odorant.

In order to investigate which ET regime is the dominated mechanism, we referred to the experimental results described by Paulson et al[66]. They performed a series of experiments to determine the rate of intramolecular charge transfer, finding that the free energy of formation (ΔG0 I) of the intermediate state was the critical variable. Specifically, the super-exchange is the dominant mechanism when ΔG0 I is large (2 eV), while the sequential mechanism will dominate when ΔG0 I is small (0.5 eV). In our DBA model, we have three relevant diabatic states denoted by D^-^BA, DB^-^A, and DBA^-^ corresponding to the initial, intermediate, and final electronic states of the system. According to Jortner’s theory [14], to undergo ET with energy conservation in the mixed quantum classical regime, the energy difference between D^-^BA and DB^-^A states, ΔG0 I, should be comparable to the sum of the reorganization energy of the environment λ_S_ and the initial product vibrational energy (υħω), otherwise the ET process becomes suppressed. Despite λ_S_ being unknown experimentally for the OR, we know that for primary ET processes in the protein photosynthetic reaction centers λs ≈ 500-3000 cm^-1^ [67]. Thus, ΔG0 I values are in the range of 10^-3^ eV–1 eV. In combination with the experimental evidences[66], we speculated that the sequential hopping regime is the dominated ET mechanism for olfaction.

The positively charged molecules may be more stable than negatively molecules, Bittner et al.[27] suggested that the charge transfer type of the DBA model is the hole transfer, and the odorant molecule undergoes electronic change from the neutral state to the cationic state. However, Pshenichnyuk et al. proves the electron-accepting properties of odorant molecules are involved in their smell recognition with experiments.[25]. Their study is based on the theory which is known as dissociative electron attachment (DEA) mechanism [68,69], using DEA spectra as odor characteristic spectra can interpret a series of mustard oil odorants structurally close odorants with markedly different odor characters.

Our DBA model agrees with the DEA mechanism in terms of resonance electron attachment with formation of the temporary negative ion and its decay by electron detachment, but it should be noted that the DEA mechanism involves an irreversible chemical reaction, i.e., dissociation of the odorant anion. We assume that most of the negatively charged odorants do not dissociate at room temperature in the air, Brookes [70] stated that olfaction does not alter the chemical composition of the odorant. To investigate whether hole transfer or electron transfer dominates the signal transduction in the OR, we also calculated hole transfer spectra for each molecule mentioned in this article with the B3LYP/6-31G method. For example, as shown in Figure 4, the calculated hole transfer spectrum of HCN has no overlap with that of benzaldehyde. We conclude that neither hole transfer absorption spectra nor emission spectra can serve as a measure of odor character.

**Figure 4.**
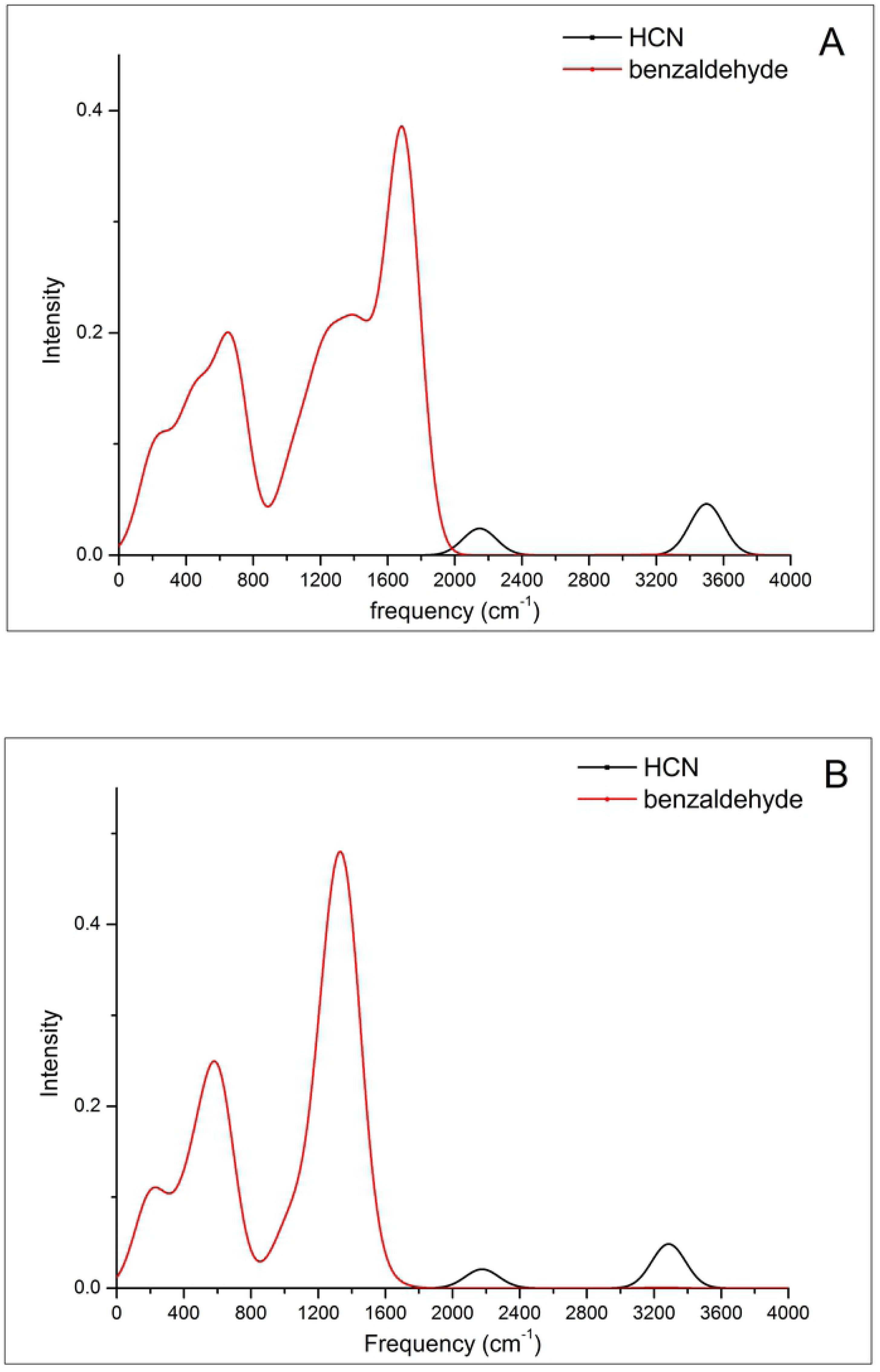
Comparative spectral analysis in hole transfer way for HCN and benzaldehyde. A. the ET absorption spectra. B. the ET emission spectra.

Block et al.[71] performed some experiments at the molecular level to compare the responses of some musk ORs to their responsive isotopomers. They determined that OR5AN1 has similar responses to its undeuterated and deuterated ligands (cyclopentadecanone and cyclopentadecanone-d28); therefore, they asserted that undeuterated and deuterated cyclopentadecanones have the same musk character and that the vibrational theory of olfaction is implausible. Turin et al.[72] suppose that the negative experimental results cannot exclude the possibility that have other ORs that are weakly activated by cyclopentadecanone or its deuterated isotopomer. Such negative evidence does not prove that the isotopomers can elicit the similar odor perception, i.e., knowing that an odorant activates a given receptor does not enable prediction of the ultimate odor percept[73]. Triller et al.[74] investigated the response of a human olfactory receptor OR1D2 to a broad array of muguet odorants. They observed that OR1D2 activation does not correlate directly with any specific odor percept, some molecules that fail to activate OR1D2 can also elicit muguet character.

Block et al.[71] also asserted that there are no experimental data showing direct evidence of electron transfer, or the effect of odorant vibrations being responsible for triggering ORs response. They further claimed that no evidence exists that GPCRs require electron transfer for their activation. One major objection to this statement is the existence of the case of the rhodopsin, a photoreceptor belonging to the GPCR class, a chromophore called 11-cis-retinal that binds to the rhodopsin via a Schiff base (RSB) and the ε-amino group of a lysine side chain in the middle of TM7[75]. In the ground state, the 11-cis-retinal is positively charged because the retinal RSB is protonated (RSBH^+^). When photoexcitation occurs, the bond of the retinal is altered, charge transfer occurs, and the positive charge is displaced from the RSBH+ to the β-ionone ring, leading to a neutralization of the RSBH^+ [75-78]^.

From the point of view of molecular interaction, if the two molecules are close enough to interact with each other, there is a charge transfer energy component in the total intermolecular interaction energy[79-81]. By the energy decomposition analysis (EDA), the intermolecular interaction energy can be partitioned into energy components such as electrostatic, polarization, charge transfer, exchange and correlation contributions and related chemical henomena[82]; here, charge transfer refers to interactions between the occupied orbital of the donor molecule and unoccupied orbital of the acceptor molecule, and the orbital energy gap and overlap are important factors. One quantum mechanics approach, Fragment Molecular Orbital (FMO), provides accurate information on the individual contribution of each residue of protein to the ligand-binding energy[83]. Heifetz et al.[84] have applied the FMO method to 18 Class A GPCR−ligand crystal structures; they observed that the average charge transfer fraction contribution for the 18 crystal structures is up to 17%.

## Acknowledgments

This work was supported by grants from the Doctoral Scientific Research Foundation of Anhui medical university (No. 0103028101). We thank the High-Performance Grid Computing Platform of Sun Yat-Sen University for supporting the quantum chemistry calculations.

